# Dynamics of microbial community composition and soil organic carbon mineralization in soil following addition of pyrogenic and fresh organic matter

**DOI:** 10.1101/033811

**Authors:** Thea Whitman, Charles Pepe-Ranney, Akio Enders, Chantal Koechli, Ashley Campbell, Daniel H. Buckley, Johannes Lehmann

## Abstract

Pyrogenic organic matter (PyOM) additions to soils can have large impacts on soil organic C (SOC) cycling. Because the soil microbial community drives SOC fluxes, understanding how PyOM additions affect soil microbes is essential to understanding how PyOM affects SOC. We studied SOC dynamics and surveyed soil microbial communities after OM additions in a field experiment. We produced and applied either 350°C corn stover PyOM or an equivalent amount of dried corn stover to a Typic Fragiudept soil. Stover increased SOC-derived and total CO_2_ fluxes (up to 6x), and caused rapid and persistent changes in bacterial community composition over 82 days. In contrast, PyOM only temporarily increased total soil CO_2_ fluxes (up to 2x) and caused fewer changes in bacterial community composition. 70% of the OTUs that increased in response to PyOM additions also responded to stover additions. These OTUs likely thrive on easily-mineralizable C that is found both in stover and, to a lesser extent, in PyOM. In contrast, we also identified unique PyOM-responders, which may respond to substrates such as polyaromatic C. In particular, members of *Gemmatimonadetes* tended to increase in relative abundance in response to PyOM but not to fresh organic matter. We identify taxa to target for future investigations of the mechanistic underpinnings of ecological phenomena associated with PyOM additions to soil.

## 1 Introduction

Large inputs of pyrogenic organic matter (PyOM) in fire-affected ecosystems can constitute up to 80% of total soil organic carbon (SOC) (Lehmann *et al*., 2008). Whether PyOM is produced naturally in fires (Czimczik and Masiello, 2007), intentionally for carbon (C) management, and/or as an agricultural amendment (Lehmann, 2007; Laird, 2008), it is important to understand how it affects the C cycle (Whitman *et al*., 2010). PyOM additions to soil can significantly affect plant growth and crop yields (*e.g*., Jeffery *et al*., 2011) and SOC dynamics (Watzinger *et al*., 2014; Maestrini *et al*., 2014; Whitman *et al*., 2015). For example, among other effects, PyOM can change soil pH (Gul *et al*., 2015), add nutrients to soils (Enders *et al*., 2012), and alter soil water-holding capacity (Abel *et al*., 2013). While PyOM additions to soils can alter inorganic C dynamics through changes to soil pH, a key mechanism for changes in SOC mineralization is altered microbial mineralization of SOC in response to PyOM (Maestrini *et al*., 2014; Whitman *et al*., 2015). Hence, an understanding of the effects of PyOM on the soil microbial community is required to understand the effects of PyOM on SOC stocks.

There are many ways PyOM could affect soil microorganisms (Kuzyakov and Bol, 2004; Lehmann *et al*., 2011; Ameloot *et al*., 2013), including, but not limited to, microbial use of PyOM as a source of energy or nutrients, changes in soil physical or chemical characteristics (*e.g*., pH), impacts on plant growth, and interference with microbial signaling (Masiello *et al*., 2013). Recent research has only begun to identify PyOM effects on soil microbial communities, and it is clear that PyOM additions to soil can induce changes in soil microbial community composition. Most current evidence has been gathered using fingerprinting approaches, such as T-RFLP (Bingeman *et al*., 1953; Jin, 2010; Kolton *et al*., 2011) or DGGE (Kolton *et al*., 2011; Chen *et al*., 2013), or by surveying PLFAs to assess microbial diversity at low phylogenetic resolution (Dunavin, 1969; Jindo *et al*., 2012; Gomez *et al*., 2014; Watzinger *et al*., 2014; Mitchell *et al*., 2015). In addition, some high-throughput DNA sequencing approaches have been applied to survey microorganisms in PyOM systems. For example, significant differences were found between soil bacterial communities in Amazonian Dark Earth soils (amended with PyOM thousands of years ago), PyOM isolated from the Amazonian Dark Earth soils, and adjacent unamended Acrisols, by surveying SSU rRNA genes (Taketani *et al*., 2013). A handful of other studies have used high-throughput sequencing approaches to characterize PyOM effects on soil microbial communities *(e.g*., Nielsen *et al*., 2014; Xu *et al*., 2014). However, due to the low number of studies and the wide diversity of PyOM materials, addition rates, soils, and environmental conditions, it is difficult to draw generalizable conclusions about the effects of PyOM on soil microbial communities, particularly at the level of individual taxa.

In this study, we investigated the effects of PyOM additions on SOC mineralization and soil microbial community composition in a field setting over 12 weeks. We also included a treatment where plots received a mass of fresh corn stover equivalent to the mass required to produce the PyOM we added. This treatment serves as a system-level control that addresses the question, “What if a given amount of biomass were not used to produce PyOM, but were applied directly to the soil?” We predicted that there would be significant differences in C dynamics between the two systems, with fresh biomass decomposing faster, and having a greater effect on SOC mineralization. Additionally, we predicted that the addition of fresh biomass would induce the greatest changes in the microbial community, with organisms that access easily-mineralizable C sources proliferating initially, and the organisms that are able to decompose aromatic or insoluble substrates, such as lignin or cellulose, emerging later. We expected that the majority of the micro-organisms that increase in relative abundance in response to PyOM additions will also increase in response to fresh organic matter additions, but they will respond to PyOM to a smaller degree, since a smaller fraction of the PyOM-C is easily-mineralizable.

## 2 Materials and methods

### 2.1 Experimental design

We conducted a field trial, with soil left unamended, soil amended with ^13^C-labelled 350°C corn stover (Zea *mays* (L.)) PyOM, or soil amended with fresh corn stover additions (Supplementary Tables 1 and 2). The ^13^C label allowed for the C sources to be partitioned between SOC and PyOM-C or corn stover-derived C (Whitman and Lehmann, 2015). The corn stover addition was designed so that the dried original corn biomass was equivalent to that which would have been required to produce the mass of corn PyOM that was applied to each plot. *(I.e*., 4.1 Mg ha^-1^ corn-derived PyOM were applied, which, with 0.365 mass fraction conserved during PyOM production, translates into 11.2 Mg ha^-1^ corn stover, which is representative of a productive corn crop in the US (Shinners and Binversie, 2007)).

The field site is located in Cornell’s research fields in Mt. Pleasant, N.Y., and is a Mardin soil (Coarse-loamy, mixed, active, mesic Typic Fragiudept) (Supplementary Table 2). The soil has been historically planted to a potato, rye, clover rotation, for the past > 30 years, but was kept in rye-clover rotation for the past 5 years, with one planting of sudangrass 3 years ago. The plot was sprayed with Roundup (glyphosate) herbicide in the fall of 2012, ploughed on May 3, 2013, and kept weed-free by hand-weeding and water-permeable landscape fabric through the summer until trial initiation.

The trial initiation date was August 16, 2013 (Day 0). Square plots (0.7 m × 0.7 m for the soil-only and PyOM, 0.45 m × 0.45 m for the stover) were surrounded by 0.7-m wide weed-free borders, maintained by hand weeding. Treatments were organized using a spatially balanced complete block design with 16 replicates of the unamended and PyOM plots, and 8 replicates of the stover plots (van Es *et al*., 2007). Soil (6.1 kg) was removed from the surface of each plot, combined with stover or PyOM additions, if needed, and mixed in a V-mixer (Twin shell dry blender, Patterson-Kelley, East Stroudsburg, PA, USA). Amended plots received 4.1 Mg ha^-1^ of PyOM or 11.2 Mg ha^-1^ of dried original corn stover. After mixing, soils were returned to their respective plots and evenly spread at the surface. Soil was gently tamped down using a flat piece of plywood. Two soil respiration collars made from 194 mm diameter white polyvinylchloride pipes were installed directly adjacent to each other at the centre of each plot with the collar protruding 30 mm and reaching 30 mm into the ground. Plots, including soil collars, were covered with water-permeable landscape fabric except during measurement for the first 2 weeks, after which fabric was removed and plots were kept weed-free by hand-weeding multiple times a week. The plots were not fertilized or watered during the trial, but were exposed to natural rainfall (temperature and precipitation are plotted in Supplementary Figure 1).

### 2.2 Biomass production

Two sets of corn plants *(Zea mays* (L.)) were grown, one in an enriched ^13^CO_2_ atmosphere growth chamber and the other in an ambient ^13^CO_2_ greenhouse. The labeled plants were grown in potting mix in a Percival AR-100L3 CO_2_-controlled growth chamber (Percival, Perry, IA). The plants were exposed to cycles of 18 h light / 6 h darkness. During light cycles, the atmosphere was maintained at 400 ppm CO_2_. During the dark cycle, CO_2_ was allowed to accumulate through respiration, and was then drawn down by photosynthesis during the next light cycle. This was done in order to reduce net respiratory losses of labeled ^13^CO_2_. Plants were pulse-labeled with 13 L of 99% ^13^CO_2_ at regular intervals over the course of their growth in order to produce an even label. Pulse labels were delivered by opening the ^13^CO_2_ cylinder to fill a balloon with ~500 mL ^13^CO_2_. The balloon remained attached to the cylinder so that the ^13^CO_2_ slowly diffused out of the balloon, delivering the pulse at a rate so that the total atmospheric concentration of CO_2_ was not affected. Plants in the growth chambers and the greenhouse were harvested just before they reached reproductive maturity and were oven-dried at 70°C.

### 2.3 PyOM production and amendment mixing

Oven-dried corn plants were ground in a hammer mill (Viking) to < 2 mm. The milled corn was pyrolyzed in a modified Fisher Scientific Isotemp programmable muffle furnace (Thermo Fisher Scientific, Waltham, MA, USA) (described in detail in Güereña *et al*. 2015) by ramping at 5°C min^−1^ to 350°C, then holding at 350°C for 45 minutes, under Ar (Supplementary Table 1). The ^13^C-labeled and natural abundance corn PyOM materials were mixed together to produce a δ^13^C value of +37.5‰. For the corn biomass-only plots, mixtures of ^13^C-labeled and natural abundance corn with a δ^13^C value of +1.7‰ were created. Mixing was done in plot-level batches to ensure that each plot received exactly these proportions of labeled and unlabeled materials.

### 2.4 CO_2_ flux and ^13^CO_2_ measurements

Soil CO_2_ flux was measured using a LI-6400XT infra-red gas analyser with a 6400–09 soil CO_2_ flux chamber attachment (LI-COR, Lincoln, Nebraska). Three flux measurements were taken in succession for each plot and averaged. Measurements were taken on days 0, 1, 2, 3, 4, 5, 6, 7, 9, 11, 12, 14, 16, 18, 26, 30, 34, 38, 41, 45, 49, 53, 57, 62, 66, 74, and 81. Additional gas samples were taken for ^13^CO_2_ analysis and emissions partitioning on day 12 and on day 66 using modified static Iso-FD chambers (Nickerson *et al*., 2013; Whitman and Lehmann, 2015) (Supplementary Figure 2).

### 2.5 Soil sampling

Soil samples for microbial community analyses were taken on days 1, 12 (chosen because it was after the first major rainfall; Supplementary Figure 1), and 82. Samples were sieved to < 2 mm, and immediately frozen in Whirl-Paks in liquid N_2_ and stored at -80°C. On days 1 and 12, two 25-mm deep soil probe samples were pooled, while on day 82, soil within the entire collar was destructively sampled.

### 2.6 Isotopic partitioning of CO_2_ samples

We partitioned the total CO_2_ fluxes between SOC and stover or PyOM-C on days 12 and 66. To determine the relative contributions of SOC and added C (either PyOM or corn stover) to soil CO_2_ fluxes, a standard isotope partitioning approach was applied (Balesdent and Mariotti, 1996). For example, for the plots with PyOM additions, the isotopic signature of the total emissions will be:

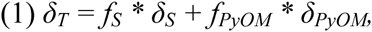

where *δ* represents the ^13^C signature of CO_2_ from total CO_2_ *(δ_T_)*, soil alone *(δ_S_)*, or PyOM *(δ_PyOM_)*, and *f_S_* and *f_PyOM_* represent the fraction of total emissions made up by soil and PyOM, respectively (Werth and Kuzyakov, 2010). Since *δ_T_, δ_S_*, and *δ_PyOM_* were all measured, and we know

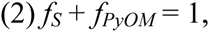

we can solve this system of two equations for the two unknown values, *f_S_ + f_PyOM_*.

### 2.7 Data processing and statistical analyses for CO_2_ fluxes

On rare occasions toward the end of the trial, we recorded negative CO_2_ fluxes, which we interpret as experimental error due to low flux rates, and have excluded these values from analyses. In addition, we excluded two data points where recorded fluxes were 56 and 16 SD away from the mean of the remaining plots. All statistical analyses were performed in R (R Core Team, 2015). CO_2_ fluxes were evaluated using a linear mixed effects model, with amendment, day, interaction between amendment and day, and plot ID (a repeated measures approach) as factors, using the R package “lme4” (Bates *et al*., 2014). To make post-hoc comparisons, we performed pairwise comparisons between the different soil amendments for a given day with a Tukey adjustment of *p*-values, using the “lsmeans” R package (Lenth, 2014).

### 2.8 Microbial community analyses

DNA was extracted from 0.25 g moist soil samples using the MoBio PowerLyzer PowerSoil kit, following the kit’s directions. The DNA was quantified using the Quant-iT PicoGreen dsDNA Assay Kit (Life Technologies) with a multimode microplate reader (Molecular Devices, Sunnyvale, CA). DNA yields were normalized on the basis of grams of dry soil extracted. SSU rRNA genes were PCR amplified in triplicate for each sample. PCR was conducted with 12.5 μL Q5 Hot Start High Fidelity 2X mastermix (New England Biolabs), 5 μL of DNA template diluted with water at a ratio of 1:50, 5 μL water, and 2.5 μL primer mixtures to a total volume of 25 μL. Each PCR consisted of a 98°C hold for 30s, followed by 30 cycles of [5s at 98°C, 20s at 20°C, and 10s at 72°C], with a final extension for 2min at 72°C. Modified 515F and 907R primers (Supplementary Tables 3 and 4) were used to target the V4/V5 regions of the 16S ribosomal RNA gene. Unique barcodes were added to the primers used for each sample so that the SSU rRNA sequences from each sample could be demultiplexed post-sequencing (Supplementary Tables 3 and 4). Each PCR product was run on a 0.5% agarose gel, along with the negative control, to determine whether amplification was successful. Replicate PCR products were pooled, and DNA concentrations were normalized across all samples using SequalPrep normalization plates (Applied Biosystems). The pooled sample was purified using a Wizard SV Gel and PCR Clean-Up System (Promega). This sample was submitted with sequencing primers (Supplementary Table 5) for paired ends 2 x 300 bp sequencing on the Illumina MiSeq v3 platform at Cornell’s Biotech Core Facility.

### 2.9 Microbial community bioinformatics

#### 2.9.1 Paired read merging, demultiplexing, and quality control

We merged the forward and reverse reads using the Paired End reAd mergeR (PEAR) (Flouri and Zhang, 2013) and demultiplexed them by barcode. DNA sequences were managed using screed databases (Nolley and Brown, 2012).

We removed merged reads with more than one expected error using USEARCH (Edgar, 2013). We also removed any sequences that had ambiguous base calls. We further identified erroneous sequences with alignment based quality control (Schloss *et al*., 2009), by aligning our sequences to the SILVA Reference Alignment as provided by the Mothur developers and removing reads that did not align to the expected region of the SSU rRNA gene. We then removed any sequences that were less than 370 bp or more than 376 bp long or had homopolymers (runs of the same base in a row) more than 8 nucleotides in length. Quality control removed 6 265 604 reads, leaving 10 237 689 high quality sequences. With 3 treatments, two with 16 replicates and one with 8 replicates, and 3 time-points, we had a total of 120 samples. Individual samples contained from 8 830 to 194 356 total sequences, with a mean of 69 419 ± 44 070.

#### 2.9.2 Operational taxonomic unit (OTU) picking

Reads were clustered into OTUs using the UPARSE methodology (Edgar, 2013) with an OTU sequence identity cut-off of 97%. In the UPARSE workflow, chimeras are detected and discarded when OTU centroids are selected. Of the quality-controlled reads, 81% could be mapped to OTU centroids. We taxonomically annotated OTU centroids using the “uclust” based taxonomic annotation framework in QIIME (v1.8) with default parameters (Caporaso *et al*., 2010; Edgar, 2013). OTUs annotated as “Archaea”, “mitochondria”, or “*Eukarya*” were removed in downstream analyses. One sample (an unamended plot from day 82) was left with only 8 sequences, so it was excluded from further analyses.

#### 2.9.3 Community analysis

OTU centroids were taxonomically annotated within the Greengenes taxonomic nomenclature using the “uclust” taxonomic annotation framework in QIIME with default parameters (Caporaso *et al*., 2010). Centroids from 97% sequence identity clusters of Greengenes database SSU rRNA gene sequences (version 13_8) and corresponding annotations were used as reference for taxonomic annotation. We used the R package “vegan” (Oksanen *et al*., 2015) to perform a nonmetric multidimensional scaling (NMDS) analysis on the weighted UniFrac distance between communities across all timepoints and amendments (Lozupone *et al*., 2011). Weighted UniFrac accounts for the relative abundance of each OTU, not just considering its presence/absence. We then tested whether there were significant effects on the microbial community due to amendment types and day of sampling with a nonparametric multivariate analysis of variance (NPMANOVA) on weighted UniFrac distances, using the “adonis" function from the R package “vegan” (Oksanen *et al*., 2015). Because we found day and amendment both had significant effects *(p* < 0.001), we performed a separate weighted UniFrac analysis and NPMANOVA for each day and amendment type (comparing each amendment to either each other or to no additions, for each day), adjusting *p*-values using a Bonferroni correction for 6 comparisons *(p_adj_* = *p* * 6).

We used the R package “DESeq2” (Love *et al*., 2014) to calculate the differential abundance (log2-fold change in relative abundance of each OTU) for each amendment type as compared to the unamended plots for both sampling days (McMurdie and Holmes, 2014). We independently filtered out OTUs that were sparsely represented across samples *(i.e*., those OTUs for which the DESeq2-normalized count across samples (“baseMean”) was less than 0.6). Sparse OTUs will not contain sufficient sequence counts to provide statistically significant results and their removal reduces the number of multiple comparisons performed, thereby mitigating problems associated with multiple comparisons to some extent. We adjusted the *p*-values with the Benjamini and Hochberg (BH) correction method and selected a study-wide false discovery rate (FDR) of 10% to denote statistical significance (Love *et al*., 2014). We defined “responding OTUs” as OTUs with a differential abundance greater than 1 and an adjusted *p*-value of <0.1. We performed a BLAST (nucleotide blast, version 2.2.29+, default parameters) search with OTU centroid sequences of responding OTUs, against the Living Tree Project (LTP) database (version 115) (Yarza 2008). The LTP database contains 16S rRNA gene sequences for all sequenced archaeal and bacterial type strains.

## 3 Results

### 3.1 Soil C dynamics

Total CO_2_ fluxes from plots that received uncharred stover additions were significantly (mixed model repeated measures design, Tukey-adjusted post-hoc comparisons, *p* < 0.05) higher than all other plots for the first 26 days and for 19 of the 27 days for which fluxes were determined (Figure 1). Stover additions had the greatest effects on CO_2_ fluxes (CO_2_ fluxes 6 times those of unamended soils) on days 7 and 11, after strong rain events (Supplementary Figure 1). Plots with PyOM additions experienced significantly higher CO_2_ fluxes than plots with no additions for the first 12 days, after which there were no significant differences (Figure 1). The increases in fluxes with PyOM additions were much less dramatic than those in the stover-amended plots, and were never greater than 2.1 times the CO_2_ emissions from unamended soils.

**Figure 1.**
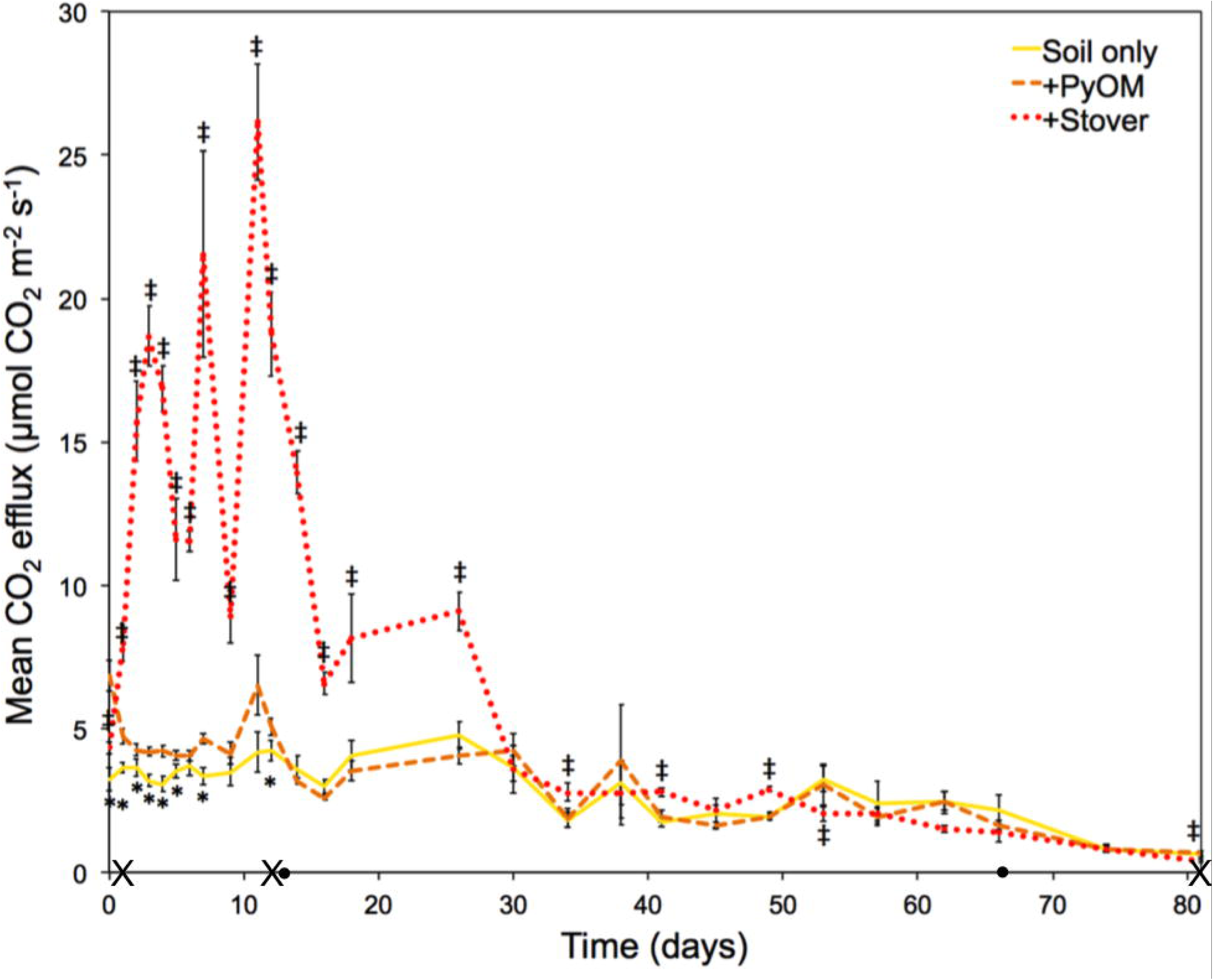
Mean CO_2_ flux rates over time. Error bars ±1 SE, n=8–16. Dotted line indicates plots that received fresh stover additions. Dashed orange line indicates plots that received PyOM additions. Solid yellow line indicates plots that had no additions. * indicates significant differences between plots with PyOM additions and stover or no-addition plots, while J indicates significant differences between plots that received fresh stover additions and PyOM or no-addition plots (mixed model repeated measures design, Tukey-adjusted post-hoc comparisons, *p* < 0.05). X indicates days where microbial community was sampled. • indicates days where ^13^CO_2_ flux was partitioned between SOC and amendments.

On day 12, ^13^C partitioning revealed that corn stover additions significantly increased SOC-derived CO_2_ fluxes as compared to soils with no additions, but PyOM additions did not (Supplementary Figure 3). On day 66, there were no significant differences in SOC-derived CO_2_ fluxes between all soils, and overall fluxes were much lower on this date (Figure 1 and Supplementary Figure 3).

### 3.2 Microbial Community Analyses

There were significant changes in the microbial community composition over time (NPMANOVA, *p* < 0.006, R^2^=0.19) and with amendment type (NPMANOVA, p<0.006, R^2^=0.20) (Figure 2). Microbial community composition in unamended plots remained relatively consistent over time (NPMANOVAs comparing Day 1 to Day 12 *[p* < 0.06, R^2^ = 0.18] and to Day 82 *[p* < 0.29, R^2^ = 0.11]), while PyOM-amended plots varied slightly and corn stover-amended plots varied greatly over time (Figure 2). The different treatments were not observed to alter microbial community composition on day 1 (only 24 hours after OM additions). By day 12, the stover treatment caused a significant change in community composition (NPMANOVA, *p* < 0.006, R^2^=0.75), though no effect of PyOM was observed. By day 82, the microbial communities in the PyOM-amended plots (NPMANOVA, *p* < 0.02, R^2^=0.19) and corn stover-amended plots (NPMANOVA, *p* < 0.02, R^2^=0.56) were both distinct from the unamended plots and from each other (NPMANOVA, *p* < 0.01, R^2^ = 0.36). We did not detect significant differences in DNA yield due to the addition of PyOM or stover directly after amendment additions (day 1), but DNA yield was significantly higher in plots with stover additions on days 12 and 82 (Supplementary Table 6).

**Figure 2.**
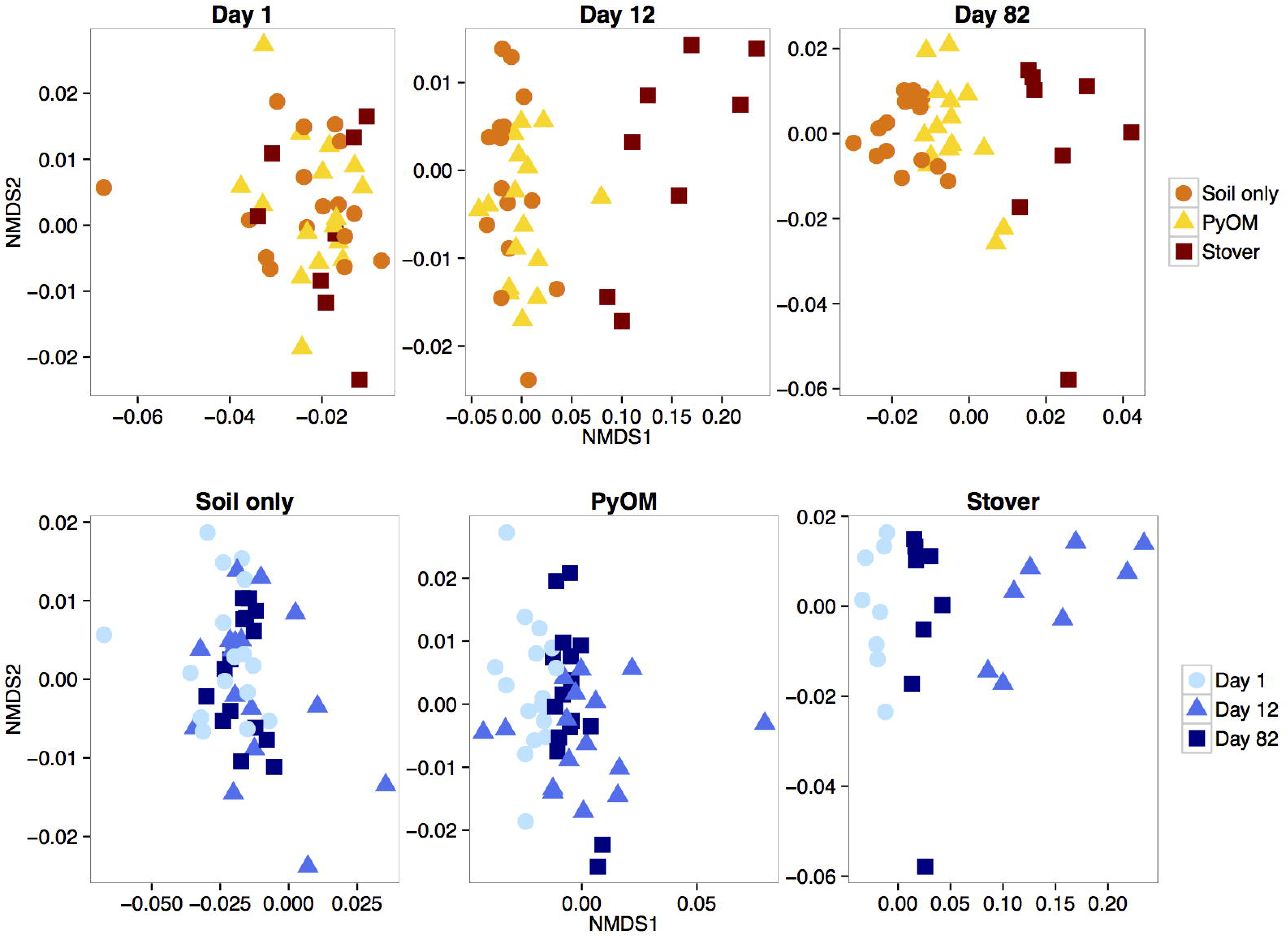
NMDS ordination (k=2, stress = 0.09) of weighted UniFrac distances between bacterial communities, showing differences across amendments for a given day (top row) and across days for a given amendment (bottom row).

The stover treatment caused multiple phyla to change significantly in relative abundance as compared to control plots on days 12 and 82 (Figure 3). In contrast, PyOM additions only led to a significant change in the relative abundance of the phyla *Armatimonadetes* (decreased), and *Bacteroidetes* (increased) and only on day 82 (Figure 3). However, we caution that decreases in relative abundance do not necessarily correspond to decreases in absolute abundance – rather, they could result from an increase in the absolute abundance of other phyla. It is notable that extracted DNA increased significantly in response to stover additions on Days 12 and 82 (Supplementary Table 6; ANOVA and Tukey’s HSD, *p* < 0.05). Since total DNA is correlated with microbial biomass (Anderson and Martens, 2013; Fornasier *et al*., 2014; Gagneux *et al*., 2011, but see Leckie *et al*., 2004), this result likely indicates an increase in microbial biomass in association with stover additions. Hence, we transformed the relative abundance data from stover treatments by scaling by the mass of extracted DNA per soil sample and performing the t-tests on those scaled values (Supplementary Note 1). The results obtained with scaled data matched those obtained with unscaled data, except declines in relative abundance for *Acidobacteria* and *Gemmatimonadetes* on day 82 were no longer statistically significant when scaled for the change in absolute DNA abundance in the stover treatment.

**Figure 3.**
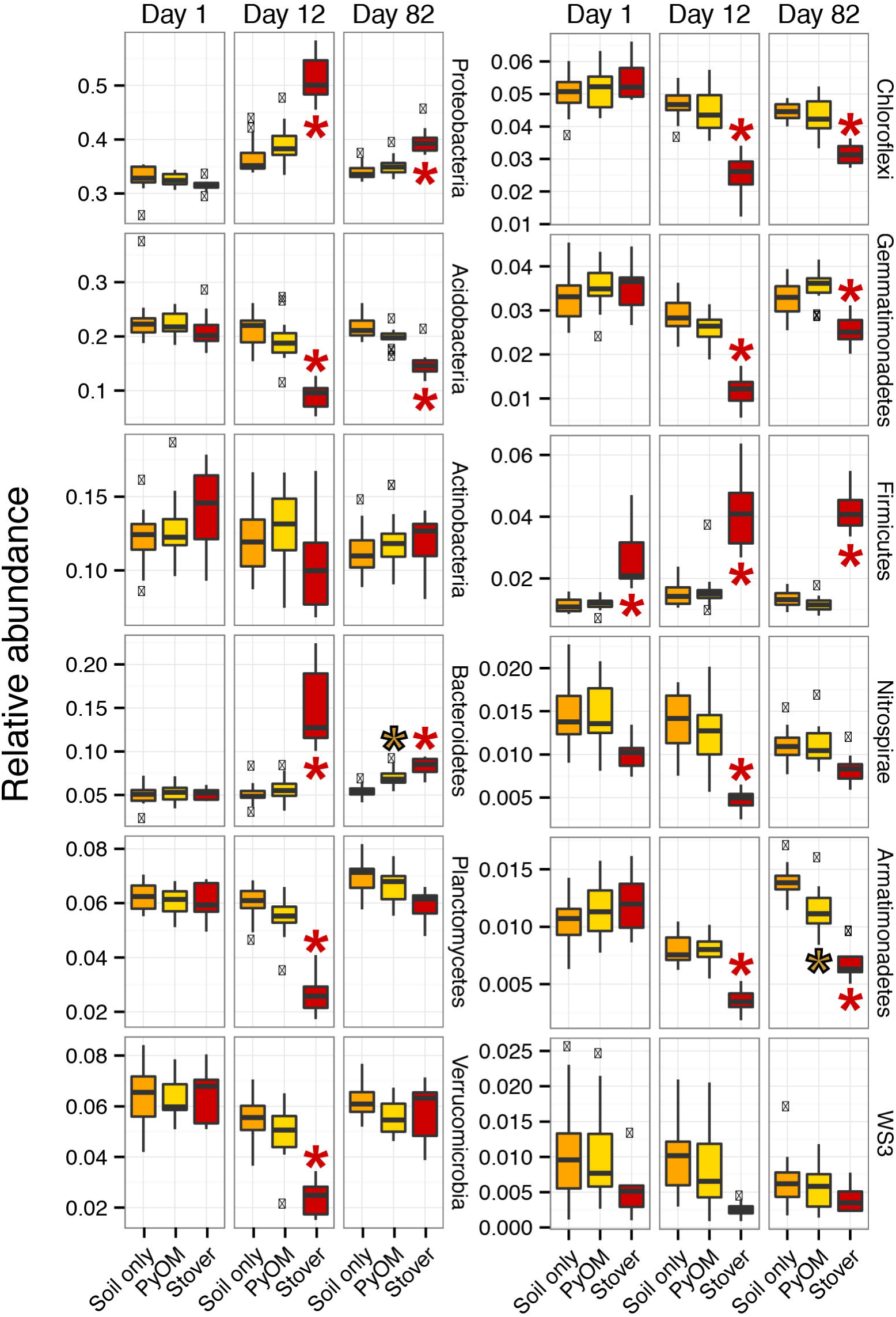
Relative abundance (unscaled by total microbial DNA) of top 12 phyla observed in the unamended, stover-amended, and PyOM-amended soils (n = 8–16). Values beyond 1.5 times the inter-quartile range are indicated (x), as are values that differ significantly from unamended (*, *t*-test, *p* < 0.05, Bonferroni-corrected for 72 comparisons).

Many OTUs, from several phyla, responded significantly to stover and/or PyOM additions (Figure 4; Supplementary Tables 7 and 8; Supplementary Figures 6–14). We use the term ‘responders’ to refer specifically to those OTUs that increase significantly in relative abundance by more than doubling in response to stover and/or PyOM additions as compared to corresponding plots that did not receive amendments. We identified 806 responders to either stover and/or PyOM from among the 7 770 total OTUs observed across all soil samples. There were more responders to stover (677 OTUs) than to PyOM (264 OTUs) (Figures 2, 4, and 5). A total of 8% of all OTUs responded specifically to stover (Figure 5, blue region), 2% responded specifically to PyOM (Figure 5, pink region), and 2% responded to both PyOM and stover (Figure 5, purple region and Supplementary Figures 4 and 5). The total number of responders to both stover and PyOM increased over time and nearly all phyla had more OTUs respond at day 82 than day 12. The only notable exception is *Firmicutes*, which had more stover responders at day 12 (36 OTUs) than day 82 (11 OTUs), and which only had 1 OTU that responded to PyOM (Supplementary Figure 6).

**Figure 4.**
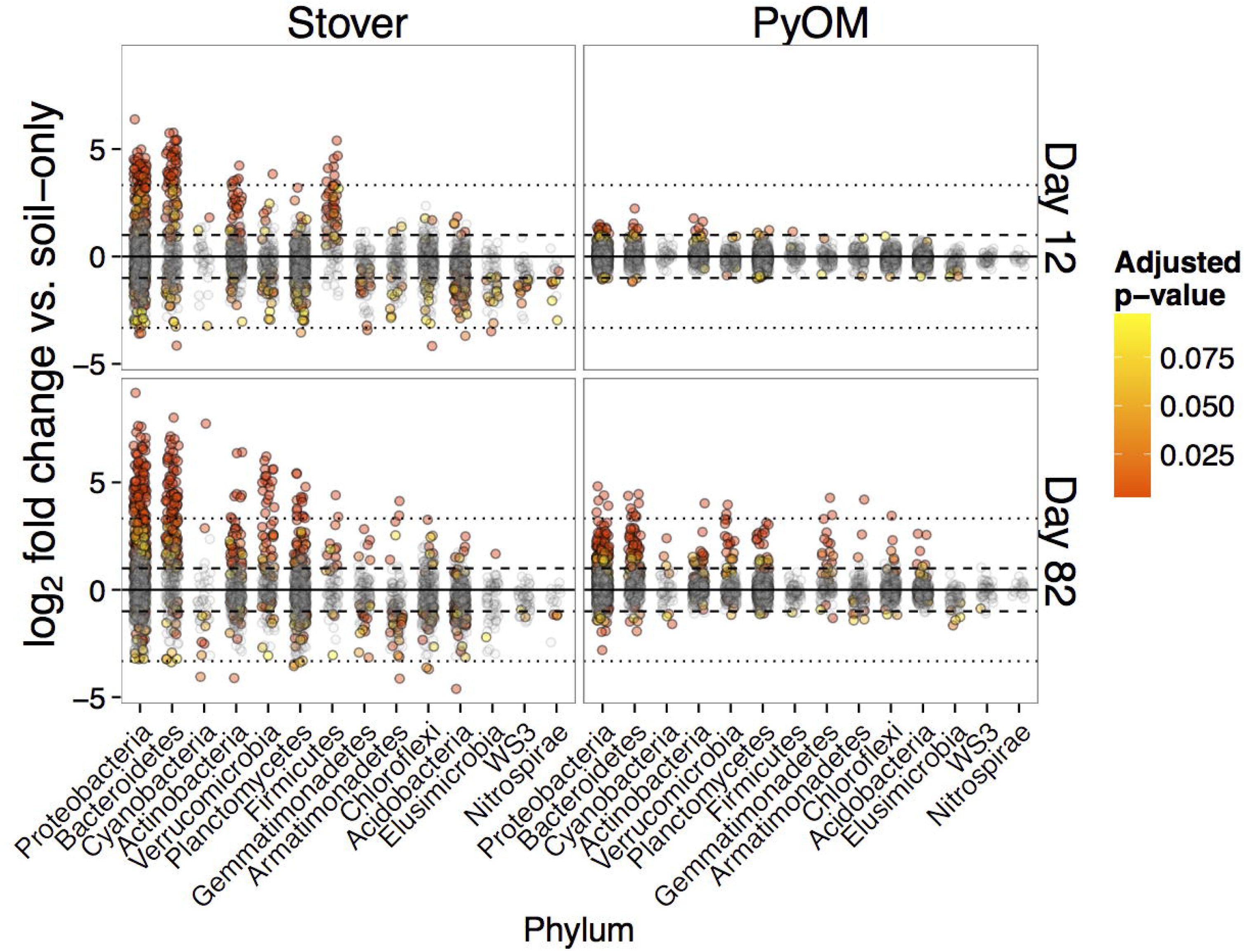
Log_2_-fold change in relative abundance of OTUs as compared to unamended plots. Each circle represents a single OTU and dashed and dotted lines represent increases or decreases of 2x and 10x, respectively. Colours are scaled from yellow to red in decreasing *p*-value, with grey points indicating OTUs with BH-adjusted *p*-values > 0. 10).

**Figure 5.**
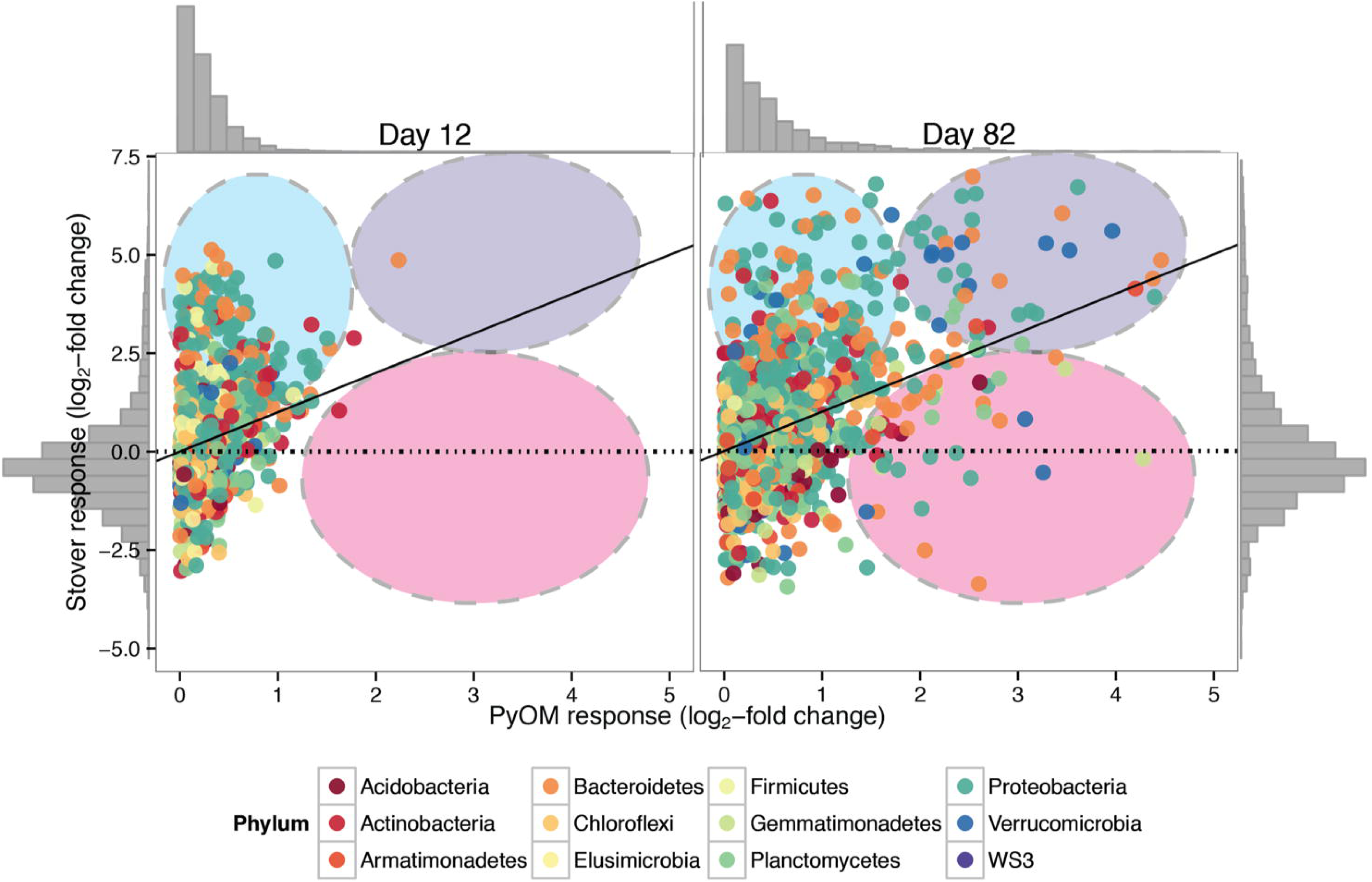
Log_2_-fold change in relative abundance of OTUs in response to stover or PyOM as compared to unamended plots. Data are the same as those depicted in Figure 4. Note different scales on axes. Each point represents the response for a single OTU across replicates, colored by phylum. Dashed oval overlays indicate response groupings, where blue indicates strong stover responders, pink indicates strong PyOM responders, and purple indicates common responders. Grey histograms at the sides of the plot indicate density of OTU points.

Many OTUs from *Firmicutes*, *Proteobacteria*, and *Bacteroidetes* responded to stover additions (Figures 4 and 5; Supplementary Figures 6, 7, and 8). Among *Proteobacteria*, 125 OTUs responded to stover additions on day 12, increasing to 235 responders on day 82 (primarily OTUs from the orders *Rhizobiales*, *Burkholderiales*, *Sphingomonadales*, *Xanthomonadales*, and *Pseudomonadales* on both days, and *Rhodospirillales* and *Myxococcales* on day 82) (Supplementary Figure 7). Among *Bacteroidetes*, 60 OTUs responded to stover additions on day 12, increasing to 128 on day 82 (primarily orders *Saprospirales, Cytophagales*, and *Sphingobacteriales*, on both days and *Flavobacteriales* on day 12) (Supplementary Figure 8). In *Firmicutes*, 36 OTUs responded to stover on day 12, declining to 11 OTUs by day 82 (orders *Bacillales* and *Clostridiales* at both times) (Supplementary Figure 6).

Only 19 OTUs were observed to respond to both stover and PyOM at day 12, but this increased to 113 OTUs by day 82. The OTUs that responded to both stover and PyOM included *Proteobacteria* (66 OTUs; Supplementary Figure 7), *Bacteroidetes* (24 OTUs; Supplementary Figure 8), and *Verrucomicrobia* (11 OTUs; Supplementary Figure 9). In particular, a number of OTUs from *Verrucomicrobia* were observed to increase greatly in relative abundance in response to both stover and PyOM as compared to untreated control plots (Figure 6 and Supplementary Figures 9 and 15).

**Figure 6.**
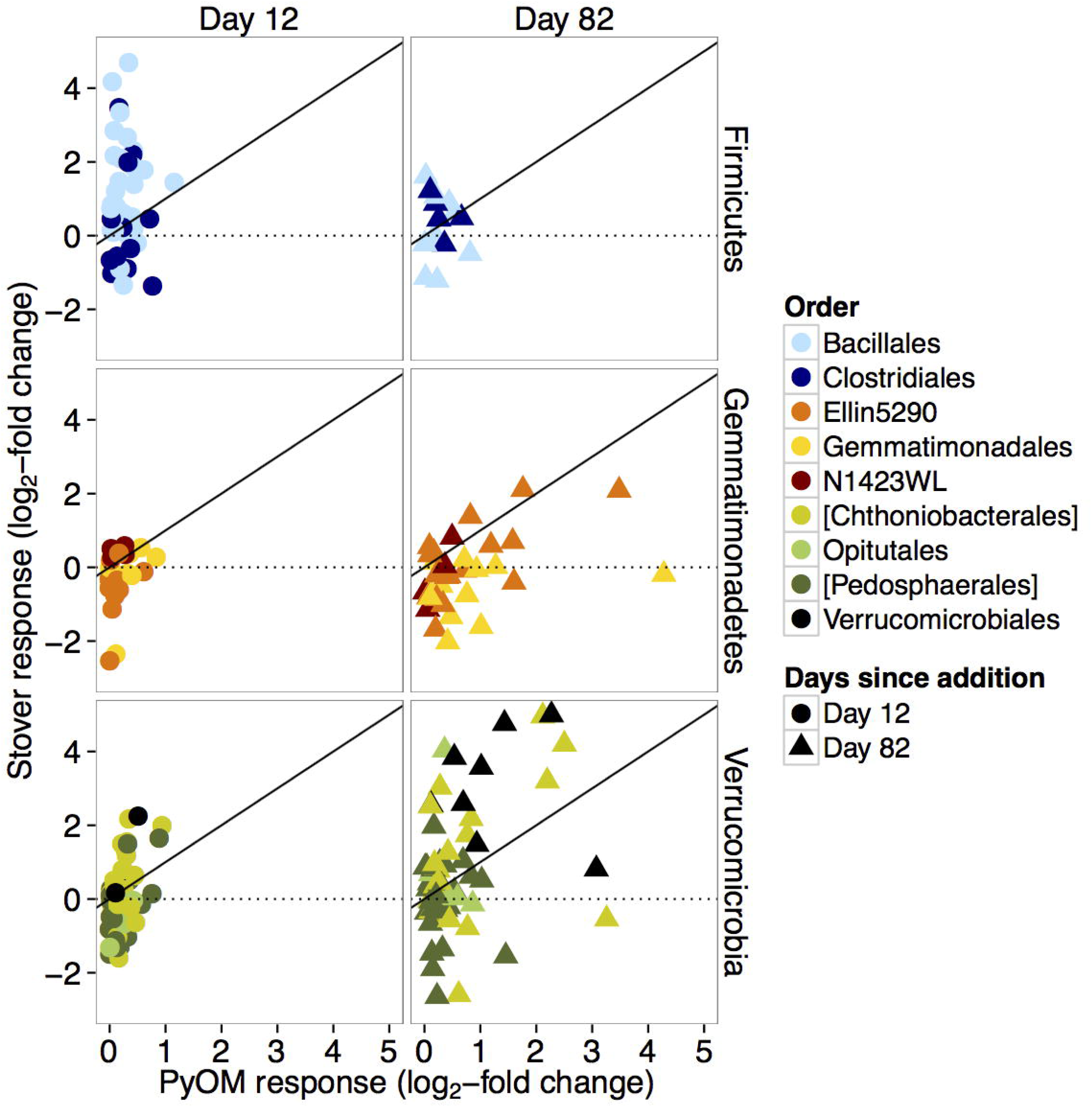
Log_2_-fold change of the relative abundance of OTUs in PyOM or stover plots *vs*. soil-only plots on days 12 and 82, for *Firmicutes, Gemmatimonadetes*, and *Verrucomicrobia* phyla. Each point represents the response for a single OTU across replicates, coloured by order. Circles represent day 12 and triangles represent day 82. Square brackets indicate candidate order in Greengenes taxonomic nomenclature.

Only 12 OTUs increased in relative abundance specifically in response to PyOM at day 12, but this increased to 138 OTUs by day 82. The OTUs that responded specifically to PyOM included *Proteobacteria* (47 OTUs; Supplementary Figure 7), *Bacteroidetes* (30 OTUs; Supplementary Figure 8), *Planctomycetes* (13 OTUs; Supplementary Figure 10), *Gemmatimonadetes* (12 OTUs; Supplementary Figure 11), and *Verrucomicrobia* (6 OTUs; Supplementary Figure 9). The strongest specific response to PyOM and not stover, by far, was observed for OTUs from *Gemmatimonadetes* (Figure 5). These included OTUs from the orders *Gemmatimonadales* and “Ellin5290” (Figure 6 and Supplementary Figure 16). Two of these *Gemmatimonadetes* OTUs were also among the top 10 most abundant PyOM responders. The most abundant PyOM responders also included two OTUs from *Bacteroidetes* (from the *Oxalobacteraceae*), and two OTUs from *Proteobacteria* (*Rhizobiales* and *Erythrobacteraceae*).

## 4 Discussion

### 4.1 Soil microbial community dynamics

The soils that received corn stover amendments showed dramatic increases in CO_2_ emissions almost immediately (Figure 1), and microbial community composition in those plots had changed significantly by day 12 (Figures 2 and 3). Stover additions resulted in significant increases in the relative abundance of OTUs from the phyla *Proteobacteria* and *Bacteroidetes*, and the orders *Actinomycetales, Bacillales*, and *Clostridiales* (Figures 3 and 4), which is consistent with previous studies (Pascault *et al*., 2013). For example, wheat residue additions to a calcareous silty clay farm soil stimulated *Firmicutes* OTUs, while alfalfa additions stimulated *Proteobacteria, Firmicutes*, and *Bacteroidetes* (Pascault *et al*., 2013). It is possible that these microorganisms are adapted to grow rapidly in response to inputs of easily-mineralizable organic matter (*e.g*., aliphatic C structures). These early-responding OTUs are likely responsible for the strong increase in total soil CO_2_ emissions during the first weeks after stover was applied (Figure 1). Since corn stover is more easily-mineralizable than pre-existing SOC, its addition might be predicted to stimulate a broad spectrum of microorganisms. However, corn stover additions stimulated only a narrow subset of microorganisms (~10% of OTUs).

Although the greatest effect of amendments on CO_2_ emissions took place within the first two weeks (Figure 1), the microbial community response to organic amendments grew stronger over 82 days. The number of PyOM responders increased 11.5-fold from day 12 to day 82, while the number of stover responders only increased 1.6-fold during this timeframe. A total of 70% of the PyOM responders that were observed by day 12 also responded to stover (Figure 5, Supplementary Figure 4). These early PyOM responders likely represent those microorganisms responsible for short-term mineralization of PyOM-C and for the CO_2_ emissions we observed within the first two weeks (although we note that this abundance-based approach would not detect possible changes to the contributions to CO_2_ efflux from microbes that did not actively grow/divide during the study (Blazewicz *et al*., 2013)). The fact that 70% of these rapidly responding PyOM responders also responded to stover suggests that they are likely metabolizing easily-mineralizable, possibly aliphatic components of PyOM which are also present within fresh corn stover (Cheng *et al*., 2008). In contrast, the OTUs that responded to both PyOM and stover on day 82 may represent microbes that were accessing the polyaromatic bulk of the PyOM-C and the remaining, less easily-mineralizable stover-C compounds (Whitman *et al*., 2013). In addition, late-responding OTUs could include microbes that are responding to other effects of the amendments, such as changes in soil physical or chemical properties, or the ecology of the system (*e.g*., changes to the soil food web, competition, or mutualisms).

### 4.2 PyOM effects on the soil microbial community

The OTUs that responded uniquely to PyOM include representatives from 14 phyla (Figure 5, pink region and Supplementary Figures 4–14). While the short-term effects of PyOM on SOC dynamics may be driven to a large extent by a relatively small, but easily-mineralizable fraction of PyOM-C (Whitman *et al*., 2014), it is essential to also understand the longer-term effects driven by chemical or physical changes to the soil environment, such as pH, soil moisture, nutrient status, or the mineralization of more chemically complex PyOM-C sources. For this, the *Gemmatimonadetes*, particularly those of classes *Gemm-5, Gemm-3*, and *Gemmatimonadales*, are a prime target (Figure 6 and Supplementary Figure 11 and 16). The first isolates of the *Gemmatimonadetes* phylum were described in 2003 (Zhang *et al*., 2003), and while little is yet known about them ecologically and physiologically, our findings seem consistent with previous observations. For example, *Gemmatimonadetes* have been found to decrease in relative abundance with the addition of wheat residues (Bernard *et al*., 2007), were more active in soil microcosms that did not receive leaf litter (Pfeiffer *et al*., 2013), and were more likely to be decomposing existing SOM than fresh OM (Pascault *et al*., 2013). Of particular relevance, *Gemmatimonadetes* increased with the addition of rice straw PyOM made at 500°C to a farmed Acrisol in a pot trial (Xu *et al*., 2014). This increase was driven by increases from the class *Gemmatimonadetes*, with some *Gemm1* and *Gemm3* increasing in relative abundance as well. These studies and our own findings suggest they may be adapted to a lifestyle associated with OM sources that are challenging to mineralize. Increased pH with PyOM additions may also have had a positive effect on *Gemmatimonadetes*, which have been reported to be more abundant in neutral pH soils (Lauber *et al*., 2009; Vishnivetskaya *et al*., 2011), although this effect was not significant for a much larger pH range than that observed in this study (2.6 vs. 0.75) (DeBruyn *et al*., 2011). *Gemmatimonadetes* may also be adapted for low soil moisture (DeBruyn *et al*., 2011), but this is not a likely explanation in our study, since we did not measure significant differences in soil moisture in the PyOM-amended soils on any sampling day (data not shown). The top 5 most abundant OTUs that responded uniquely to PyOM, including members of the *Oxalobacteraceae, Rhizobiales*, and *Erythrobacteraceae*, could also be good targets for future investigations into microbial interactions with PyOM.

PyOM produced from different materials under different conditions can result in a wide range of pH values (Enders *et al*., 2012), and in this study, PyOM additions significantly increased soil pH (Supplementary Figure 17), albeit by less than a full pH unit. Soil pH is strongly correlated with community composition (Lauber *et al*., 2009; Rousk *et al*., 2010). In particular, the aptly named phylum *Acidobacteria* has been shown to be especially sensitive to pH shifts, although its subgroups show variable responses to acidity: subgroups 1, 2, and 3 have been shown to increase at lower pHs, while subgroups 4, 5, 6, 7, and 17 have been shown to increase at higher pHs (Rousk *et al*., 2010; Bartram *et al*., 2013). Both Bartram *et al*. (2013) and Rousk *et al*. (2010) characterized soils from long-term (50+ and 100+ years, respectively) liming trials, so it is not possible to predict from those studies the expected timescale of a soil microbial community response to pH changes. However, we may ask whether the PyOM-specific response in this study is driven by pH shifts. We found that OTUs from subgroup 4 did generally increase with PyOM additions (Supplementary Figure 18). However, we also found that members of subgroup 6 decreased in relative abundance, while subgroup 3 increased with PyOM additions (Supplementary Figure 18), which is counter to what trends in previous studies would predict if these changes were being driven purely by the pH increase with PyOM additions. This does not necessarily contrast with the previous studies – rather, it may show that factors other than pH are likely important for driving the observed changes in subgroups 3 and 6 in this system. Explanations besides pH are particularly likely, as the pH shift by +0.75 units was small in comparison to the potential magnitude of natural pH gradients common in soils (*e.g*., due to biological activity, rhizosphere effects, or wet-dry cycles (Husson, 2012)). *Acidobacteria* have sometimes been characterized as being poorly equipped to compete in high nutrient conditions (Fierer *et al*., 2007). If some *Acidobacteria* are able to mineralize challenging C forms, such as fused aromatic C ring structures, but are poorly adapted for metabolizing easily-mineralizable C, for example, this could explain the positive response of OTUs in *Acidobacteria* subgroup 3 to PyOM additions on day 82, despite the accompanying small pH increase.

### 4.3 Soil C dynamics

Stover amendments had a larger effect than PyOM on both soil microbial community composition and also the mineralization of existing SOC. Stover additions significantly increased plot-level CO_2_ emissions over a longer period of time (Figure 1), and also resulted in significantly greater SOC-derived CO_2_ emissions on Day 12 (Supplementary Figure 3). However, this increase in mineralization of existing SOC disappeared by Day 66 (Supplementary Figure 3). These dynamics likely reflect the relative microbial accessibility or solubility of the two amendments (Lehmann and Kleber, 2015).

Because the PyOM was produced at a relatively low temperature (350°C), there was likely a substantial fraction of relatively easily-mineralizable C (Whitman *et al*., 2013; Zimmerman, 2010) that contributed to the increases in total CO_2_ during the first week after application (Figure 1 and Supplementary Figure 19). This fraction could include aliphatic C compounds, carboxylic acids, cellulose, and hemicellulose (Whitman *et al*., 2013). (It is also possible that a small portion of the PyOM-C losses were due to the dissolution of carbonates (Supplementary Figure 17).) However, we did not find evidence that increased microbial activity due to PyOM additions affected SOC-derived CO_2_ emissions (Supplementary Figure 3). This is likely because the PyOM-C was more challenging for microorganisms to mineralize than the stover-C, resulting in slower mineralization of PyOM as compared to the stover. While there was 1.8 times as much total stover-C added as PyOM-C, CO_2_ emissions increased with stover additions much more than 1.8 times the amount they increased with PyOM additions (Figure 1; Supplementary Figure 19). Only the stover provided a large easily-mineralizable C subsidy to the microbial community, which stimulated the microbial community, increased microbial biomass, and may have increased general enzymatic activity. This, in turn, increased mineralization of existing SOC (Blagodatskaya and Kuzyakov, 2008). The lack of any detectable net effect on SOC mineralization from PyOM additions (often termed a “priming effect” (Bingeman *et al*., 1953; Whitman *et al*., 2014; Woolf and Lehmann, 2012)) is not unexpected. Specific combinations of PyOM materials and soils have been found to produce a wide range of effects on SOC mineralization (Whitman *et al*., 2015; Maestrini *et al*., 2014). The finding that PyOM application may result in less SOC loss than the application of an equivalent amount of fresh stover has implications for residue and C management at the farm scale. In systems where organic matter sources will rapidly mineralize under baseline conditions, PyOM production from this material and its return to the soil may result in greater net C gains than adding the fresh organic matter directly to the soil, due to the strong impact of fresh organic matter on the soil microbial community and their C mineralization activity.

## Acknowledgements

We are grateful for the financial support by awards from the Towards Sustainability Foundation, Cornell Sigma Xi, NSERC PGS-D, NSF-BREAD (IOS-0965336), Cornell Biogeochemistry program, Cornell Crop and Soil Science Department, USDA-NIFA Carbon Cycle (2014-67003-22069), and the Cornell Atkinson Center for a Sustainable Future. Thanks to Nick Nickerson and Forerunner Research / EOSENSE for advice regarding chamber design. Thanks to Kelly Hanley, Joseph Amsili, Angela Possinger, Romy Zyngier, and Andy Cavin for help with fieldwork. Thanks also to Brent Whitman and Ellen Whitman for helping with lab analyses. Thanks to Dr. Christine Goodale for helpful discussion and suggestions. The authors declare no conflict of interest.

## Author contributions

T.W., D.H.B., and J.L. designed the experiment, T.W. and A.E. grew the biomass and produced the PyOM, A.E. designed and built the CO_2_ chambers, T.W. conducted the field trial, T.W. performed the biogeochemical lab work, A.C., C.K., and T.W. performed the molecular lab work, A.C., C.K., and C.P.-R. developed the bioinformatics analysis pipeline, T.W. and C.P.R. performed the bioinformatics analyses, T.W., C.P.-R., D.H.B., and J.L. interpreted the data, and T.W. wrote the manuscript, and all authors commented on the paper.

## Supplementary information

Supplementary information is available at ISME J’s website. All scripts used for microbial sequence data processing and analysis are openly available at https://github.com/TheaWhitman/PyOM. Sequence data are deposited in the NCBI Sequence Read Archive under accession number XXX.

## REFERENCES

Abel S, Peters A, Trinks S, Schonsky H, Facklam M. (2013). Impact of biochar and hydrochar addition on water retention and water repellency of sandy soil. Geoderma 202–203: 183–191.

Ameloot N, Graber ER, Verheijen FGA, De Neve S. (2013). Interactions between biochar stability and soil organisms: review and research needs. European Journal of Soil Science 64: 379–390.

Anderson T-H, Martens R. (2013). DNA determinations during growth of soil microbial biomass. Soil Biology and Biochemistry 57: 487–495.

Balesdent J, Mariotti A. (1996). Measurement of soil organic matter turnover using ^13^C natural abundance. In Mass Spectrometry of Soils, Boutton, TW & Yamasaki, SI (Eds.) Marcel Dekker, New York.

Bartram AK, Jiang X, Lynch MD, Masella AP, Nicol GW, Dushoff J, et al. (2013). Exploring links between pH and bacterial community composition in soils from the Craibstone Experimental Farm. FEMS Microbiology Ecology 87: 403–415.

Bernard L, Mougel C, Maron P-A, Nowak V, Lévêque J, Henault C, et al. (2007). Dynamics and identification of soil microbial populations actively assimilating carbon from ^13^C-labelled wheat residue as estimated by DNA- and RNA-SIP techniques. Environ Microbiol 9: 752–764.

Bingeman CW, Varner JE, Martin WP. (1953). The effect of the addition of organic materials on the decomposition of an organic soil. Soil Science Society of America Journal 17: 34–38.

Blagodatskaya E, Kuzyakov Y. (2008). Mechanisms of real and apparent priming effects and their dependence on soil microbial biomass and community structure: critical review. Biology and Fertility of Soils 45: 115–131.

Blazewicz SJ, Barnard RL, Daly RA, Firestone MK. (2013). Evaluating rRNA as an indicator of microbial activity in environmental communities: limitations and uses. The ISME Journal 7: 2061–2068.

Caporaso JG, Kuczynski J, Stombaugh J, Bittinger K, Bushman FD, Costello EK, et al. (2010). QIIME allows analysis of high-throughput community sequencing data. Nature Methods 7: 335–336.

Chen J, Liu X, Zheng J, Zhang B, Lu H, Chi Z, et al. (2013). Biochar soil amendment increased bacterial but decreased fungal gene abundance with shifts in community structure in a slightly acid rice paddy from Southwest China. Applied Soil Ecology 71: 33–44.

Cheng C-H, Lehmann J, Engelhard MH. (2008). Natural oxidation of black carbon in soils: Changes in molecular form and surface charge along a climosequence. Geochimica Et Cosmochimica Acta 72: 1598–1610.

Czimczik CI, Masiello CA. (2007). Controls on black carbon storage in soils. Global Biogeochemical Cycles 21:GB3005.

DeBruyn JM, Nixon LT, Fawaz MN, Johnson AM, Radosevich M. (2011). Global biogeography and quantitative seasonal dynamics of *Gemmatimonadetes* in soil. Applied and Environmental Microbiology 77: 6295–6300.

Dunavin LS. (1969). A comparison of Gahi-1 millet and Grazer A sorghum x sudangrass at several pH levels. Proceedings of the Soil and Crop Science Society of Florida 29: 163–168.

Edgar RC. (2013). UPARSE: highly accurate OTU sequences from microbial amplicon reads. Nature Methods 10: 996–998.

Enders A, Hanley K, Whitman T, Joseph S, Lehmann J. (2012). Characterization of biochars to evaluate recalcitrance and agronomic performance. Bioresource Technology 114: 644–653.

Fierer N, Bradford MA, Jackson RB. (2007). Toward an ecological classification of soil bacteria. Ecology 88: 1354–1364.

Fornasier F, Ascher J, Ceccherini MT, Tomat E, Pietramellara G. (2014) A simplified rapid, low-cost and versatile DNA-based assessment of soil microbial biomass. Ecological Indicators 45: 75–82.

Gagneux C, Akpa-Vinceslas M, Sauvage H, Desaire S, Houot S, Laval K. (2011) Fungal, bacterial and plant dsDNA contributes to soil total DNA extracted from silty soils under different farming practices: Relationships with chloroform-labile carbon. Soil Biology and Biochemistry 43: 431–437.

Gomez JD, Denef K, Stewart CE, Zheng J, Cotrufo MF. (2014). Biochar addition rate influences soil microbial abundance and activity in temperate soils. European Journal of Soil Science 65:28–39.

Gul S, Whalen JK, Thomas BW, Sachdeva V, Deng H. (2015). Physico-chemical properties and microbial responses in biochar-amended soils: Mechanisms and future directions. Agriculture, Ecosystems and Environment 206: 46–59.

Guereña DT, Lehmann J, Thies JE, Enders A, Karanja N, Neufeldt H. (2015) Partitioning the contributions of biochar properties to enhanced biological nitrogen fixation in common bean *(Phaseolus vulgaris)*. Biology and Fertility of Soils 51: 479–491.

Husson, O. (2012). Redox potential (Eh) and pH as drivers of soil/plant/microorganism systems: a transdisciplinary overview pointing to integrative opportunities for agronomy. Plant and Soil 362: 389–417.

Jeffery S, Verheijen FGA, van der Velde M, Bastos AC. (2011). A quantitative review of the effects of biochar application to soils on crop productivity using meta-analysis. Agriculture, Ecosystems and Environment 144: 175–187.

Jin H. (2010). Characterization of microbial life colonizing biochar and biochar-amended soils. Ph.D. dissertation Cornell University: Ithaca, NY.

Jindo K, Sanchez-Monedero MA, Hernández T, García C, Furukawa T, Matsumoto K, et al. (2012). Biochar influences the microbial community structure during manure composting with agricultural wastes. Science of The Total Environment 416: 476–481.

Kolton M, Meller Harel Y, Pasternak Z, Graber ER, Elad Y, Cytryn E. (2011). Impact of biochar application to soil on the root-associated bacterial community structure of fully developed greenhouse pepper plants. Applied and Environmental Microbiology 77: 4924–4930.

Kuzyakov Y, Bol R. (2004). Using natural ^13^C abundances to differentiate between three CO_2_ sources during incubation of a grassland soil amended with slurry and sugar. Journal of Plant Nutrition and Soil Science 167: 669–677.

Laird DA. (2008). The charcoal vision: A win–win–win scenario for simultaneously producing bioenergy, permanently sequestering carbon, while improving soil and water quality. Agronomy Journal 100: 178–181.

Lauber CL, Hamady M, Knight R, Fierer N. (2009). Pyrosequencing-based assessment of soil pH as a predictor of soil bacterial community structure at the continental scale. Applied and Environmental Microbiology 75: 5111–5120.

Leckie SE, Prescott CE, Grayston SJ, Neufeld JD, Mohn WW. (2004) Comparison of chloroform fumigation-extraction, phospholipid fatty acid, and DNA methods to determine microbial biomass in forest humus. Soil Biology and Biochemistry 36: 529–532.

Lehmann J. (2007). A handful of carbon. Nature 447: 143–144.

Lehmann J, Kleber M. (2015) The contentious nature of soil organic matter. Nature 528: 60–68.

Lehmann J, Rillig MC, Thies J, Masiello CA, Hockaday WC, Crowley D. (2011). Biochar effects on soil biota - A review. Soil Biology and Biochemistry 43: 1812–1836.

Lehmann J, Skjemstad J, Sohi S, Carter J, Barson M, Falloon P, et al. (2008). Australian climate-carbon cycle feedback reduced by soil black carbon. Nature Geoscience 1: 832–835.

Lozupone C, Lladser ME, Knights D, Stombaugh J, Knight R. (2011). UniFrac: an effective distance metric for microbial community comparison. The ISME Journal 5: 169–172.

Maestrini B, Nannipieri P, Abiven S. (2014). A meta-analysis on pyrogenic organic matter induced priming effect. GCB Bioenergy 7: 577–590.

Major J, Lehmann J, Rondon M, Goodale C. (2010). Fate of soil-applied black carbon: downward migration, leaching and soil respiration. Global Change Biology 16: 1366–1379.

Masiello CA, Chen Y, Gao X, Liu S, Cheng H-Y, Bennett MR, et al. (2013). Biochar and microbial signaling: production conditions determine effects on microbial communication. Environmental science & Technology 47: 11496–11503.

McMurdie PJ and Holmes S. (2014) Waste not, want not: Why rarefying microbiome data is inadmissible. PLoS Computational Biology 10:e1003531.

Mitchell PJ, Simpson AJ, Soong R, Simpson MJ. (2015). Shifts in microbial community and water-extractable organic matter composition with biochar amendment in a temperate forest soil. Soil Biology and Biochemistry 81: 244–254.

Nickerson N, Egan J, Risk D. (2013). Iso-FD: A novel method for measuring the isotopic signature of surface flux. Soil Biology and Biochemistry 62: 99–106.

Nielsen S, Minchin T, Kimber S, van Zwieten L, Gilbert J, Munroe P, et al. (2014). Comparative analysis of the microbial communities in agricultural soil amended with enhanced biochars or traditional fertilisers. Agriculture, Ecosystems and Environment 191: 73–82.

Oksanen J, Blanchet FG, Kindt R, Legendre P, Minchin PR, OHara RB, et al. (2015). vegan: Community Ecology Package. R package version 2.3-0. http://CRAN.R-project.org/package=vegan.

Pascault N, Ranjard L, Kaisermann A, Bachar D, Christen R, Terrat S, et al. (2013). Stimulation of different functional groups of bacteria by various plant residues as a driver of soil priming effect. Ecosystems 16: 810–822.

Pfeiffer B, Fender A-C, Lasota S, Hertel D, Jungkunst HF, Daniel R. (2013). Leaf litter is the main driver for changes in bacterial community structures in the rhizosphere of ash and beech. Applied Soil Ecology 72: 150–160.

R Core Team (2015). R: A language and environment for statistical computing. R Foundation for Statistical Computing, Vienna, Austria. URL http://www.R-project.org/.

Rousk J, Bååth E, Brookes PC, Lauber CL, Lozupone C, Caporaso JG, et al. (2010). Soil bacterial and fungal communities across a pH gradient in an arable soil. The ISME Journal 4: 1340–1351.

Schloss PD, Westcott SL, Ryabin T, Hall JR, Hartmann M, Hollister EB, et al. (2009). Introducing mothur: Open-source, platform-independent, community-supported software for describing and comparing microbial communities. Applied and Environmental Microbiology 75: 7537–7541.

Shinners KJ, Binversie BN. (2007). Fractional yield and moisture of corn stover biomass produced in the Northern US Corn Belt. Biomass and Bioenergy 31: 576–584.

Slavich PG, Sinclair K, Morris SG, Kimber SWL, Downie A, Zwieten L. (2013). Contrasting effects of manure and green waste biochars on the properties of an acidic ferralsol and productivity of a subtropical pasture. Plant and Soil 366: 213–227.

Taketani RG, Lima AB, Conceição Jesus E, Teixeira WG, Tiedje JM, Tsai SM. (2013). Bacterial community composition of anthropogenic biochar and Amazonian anthrosols assessed by 16S rRNA gene 454 pyrosequencing. Antonie van Leeuwenhoek 104: 233–242.

van Es HM, Gomes CP, Sellmann M, van Es CL. (2007). Spatially-balanced complete block designs for field experiments. Geoderma 140: 346–352.

Vishnivetskaya TA, Mosher JJ, Palumbo AV, Yang ZK, Podar M, Brown SD, et al. (2011). Mercury and Other Heavy Metals Influence Bacterial Community Structure in Contaminated Tennessee Streams. Applied and Environmental Microbiology 77: 302–311.

Watzinger A, Feichtmair S, Kitzler B, Zehetner F, Kloss S, Wimmer B, et al. (2014). Soil microbial communities responded to biochar application in temperate soils and slowly metabolized ^13^C-labelled biochar as revealed by ^13^C PLFA analyses: results from a short-term incubation and pot experiment. European Journal of Soil Science 65: 40–51.

Werth M, Kuzyakov Y. (2010). ^13^C fractionation at the root-microorganisms-soil interface: A review and outlook for partitioning studies. Soil Biology and Biochemistry 42: 1372–1384.

Whitman T, Enders A, Lehmann J. (2014). Pyrogenic carbon additions to soil counteract positive priming of soil carbon mineralization by plants. Soil Biology and Biochemistry 73: 33–41.

Whitman T, Hanley K, Enders A, Lehmann J. (2013). Predicting pyrogenic organic matter mineralization from its initial properties and implications for carbon management. Organic Geochemistry 64: 76–83.

Whitman T, Lehmann J. (2015). A dual-isotope approach to allow conclusive partitioning between three sources. Nature Communications 6:no.8708.

Whitman T, Singh BP, Zimmerman AR. (2015). Priming effects in biochar-amended soils: Implications of biochar-soil organic matter interactions for carbon storage. In: Biochar for Environmental Management, Lehmann, J & Joseph, S (eds). Routledge, NY.

Whitman T, Zhu Z, Lehmann J. (2014). Carbon mineralizability determines interactive effects on mineralization of pyrogenic organic matter and soil organic carbon. Environmental Science & Technology 48: 13727–13734.

Woolf D, Lehmann J. (2012). Modelling the long-term response to positive and negative priming of soil organic carbon by black carbon. Biogeochemistry, 111: 83–95.

Xu H-J, Wang X-H, Li H, Yao H-Y, Su J-Q, Zhu Y-G. (2014). Biochar impacts soil microbial community composition and nitrogen cycling in an acidic soil planted with rape. Environmental science & Technology 48: 9391–9399.

Yarza P, Richter M, Peplies J, Euzeby J, Amann R, Schleifer K-H, et al. (2008). The All-Species Living Tree project: A 16S rRNA-based phylogenetic tree of all sequenced type strains. Systematic and Applied Microbiology 31: 241–250.

Zhang H, Sekiguchi Y, Hanada S, Hugenholtz P, Kim H, Kamagata Y, et al. (2003). *Gemmatimonas aurantiaca* gen. nov., sp nov., a gram-negative, aerobic, polyphosphate-accumulating micro-organism, the first cultured representative of the new bacterial phylum *Gemmatimonadetes* phyl. nov. International Journal of Systematic and Evolutionary Microbiology 53: 1155–1163.

Zimmerman AR. (2010). Abiotic and microbial oxidation of laboratory-produced black carbon (biochar). Environmental Science & Technology 44: 1295–1301.

